# Scalable and Automated CRISPR-Based Strain Engineering Using Droplet Microfluidics

**DOI:** 10.1101/2021.06.10.447396

**Authors:** Kosuke Iwai, Maren Wehrs, Megan Garber, Jess Sustarich, Lauren Washburn, Zachary Costello, Peter W. Kim, David Ando, William R. Gaillard, Nathan J. Hillson, Paul D. Adams, Aindrila Mukhopadhyay, Hector Garcia Martin, Anup K. Singh

## Abstract

We present a droplet-based microfluidic system that enables CRISPR-based gene editing and high-throughput screening on chip. The microfluidic device contains a 10 x 10 element array, each element containing sets of electrodes for two electric field actuated operations-electrowetting for merging droplets to mix reagents and electroporation for transformation. It can perform up to 100 genetic modifications in parallel, providing a scalable platform for generating the large number of engineered strains required for combinatorial optimization of genetic pathways and predictable bioengineering. We demonstrate the system’s capabilities through CRISPR-based engineering of two test cases-1) disruption of the function of enzyme galactokinase (*galK*) in *E. coli* and 2) targeted engineering of glutamine synthetase gene (*glnA*) and blue-pigment synthetase (*bpsA*) enzyme to improve indigoidine production in *E. coli*.

## Introduction

The CRISPR/Cas9 (clustered regularly interspaced short palindromic repeats and its associated protein, Cas9) system has proven to be a powerful tool for genome engineering in organisms, both eukaryotes and prokaryotes. Recombineering is a method of genetic engineering relying on short 50-base pair homologous recombination. It allows insertion, deletion, or alteration of any sequence precisely and is not dependent on the location of restriction sites that typically limit genetic engineering in bacterial systems such as *E. coli*. Multiplex Automated Genome Engineering (MAGE) was developed to simultaneously introduce many chromosomal changes in a combinatorial fashion across a population of cells^1^. In *E. coli*, the CRISPR-Cas9 system has been recently coupled to λ Red oligos recombineering in order to improve its efficiency^2^. CRISPR-MAGE exploits intrinsic negative selection against the wild type of CRISPR/Cas9 in order to increase the MAGE performance for small genome modifications such as codon substitution or translation control elements^3^. The system is based on two curable plasmids that encode optimized versions of both systems: λ Red recombineering and CRISPR/Cas9. While realization of the high-throughput engineering potential of MAGE and CRISPR-MAGE require automated instrumentations, robotic workstations that integrate the requisite unit operations are expensive and not accessible to most researchers.

Using microscopic droplets as reaction chambers has proven to be a powerful approach to improve the throughput of synthetic biology experiments^4^. The benefits of this approach include faster reactions because of the small dimensions, low reagent consumption (enabling more reactions), and better control of experimental design^5^. A number of microfluidic systems have been developed, including flow-based droplet microfluidics that utilizes pressure-driven flow and electrical-based digital microfluidics (DMF) that utilizes electrowetting-on-dielectric (EWOD)^4,6–10^.

Here, we present a microfluidic platform for miniaturization and automation of CRISPR-MAGE. The device has 100 wells, bottom of each containing a set of electrodes for carrying out two functions-electrowetting based merger of droplets and electroporation for transformation of cells (**Fig. 1**). The configuration of the chip uses a 384-well template and is easily integratable with liquid handling robots (*e*.*g*., Labcyte Echo, iDOT, Hamilton, Tecan, OpenTrons, etc.), for automated sample input. The microfluidic chip is made with commonly used processes and materials, making it easier for adoption by non-microfluidic experts. The microfluidic chip was used for performing targeted genomic changes in *E. coli* through CRISPR-based MAGE (CRMAGE^3^) recombineering in an automated fashion for two test cases. First was to disrupt the function of enzyme galactokinase (*galK*) to demonstrate CRMAGE. We then targeted engineering of glutamine synthetase gene (*glnA*) and blue-pigment synthetase (*bpsA*) enzyme to improve indigoidine production. Indigoidine is a non-ribosomal peptide with potential applications as a dye, antioxidant and antimicrobial compound. It is a viable alternative to indigo, currently one of the most common blue dyes used in the fabric industry. Indigo is derived from its precursor indican (present in plant leaves), but its production involves harsh reagents and environmentally detrimental waste streams. Indigoidine provides a potential alternative through the use of environmentally benign routes in engineered bacteria and yeast^11–15^. High throughput approaches for biosynthetic pathway bioengineering to improve its productivity will be critical to generate final strains that meet industry production standards.

**Figure 1:**
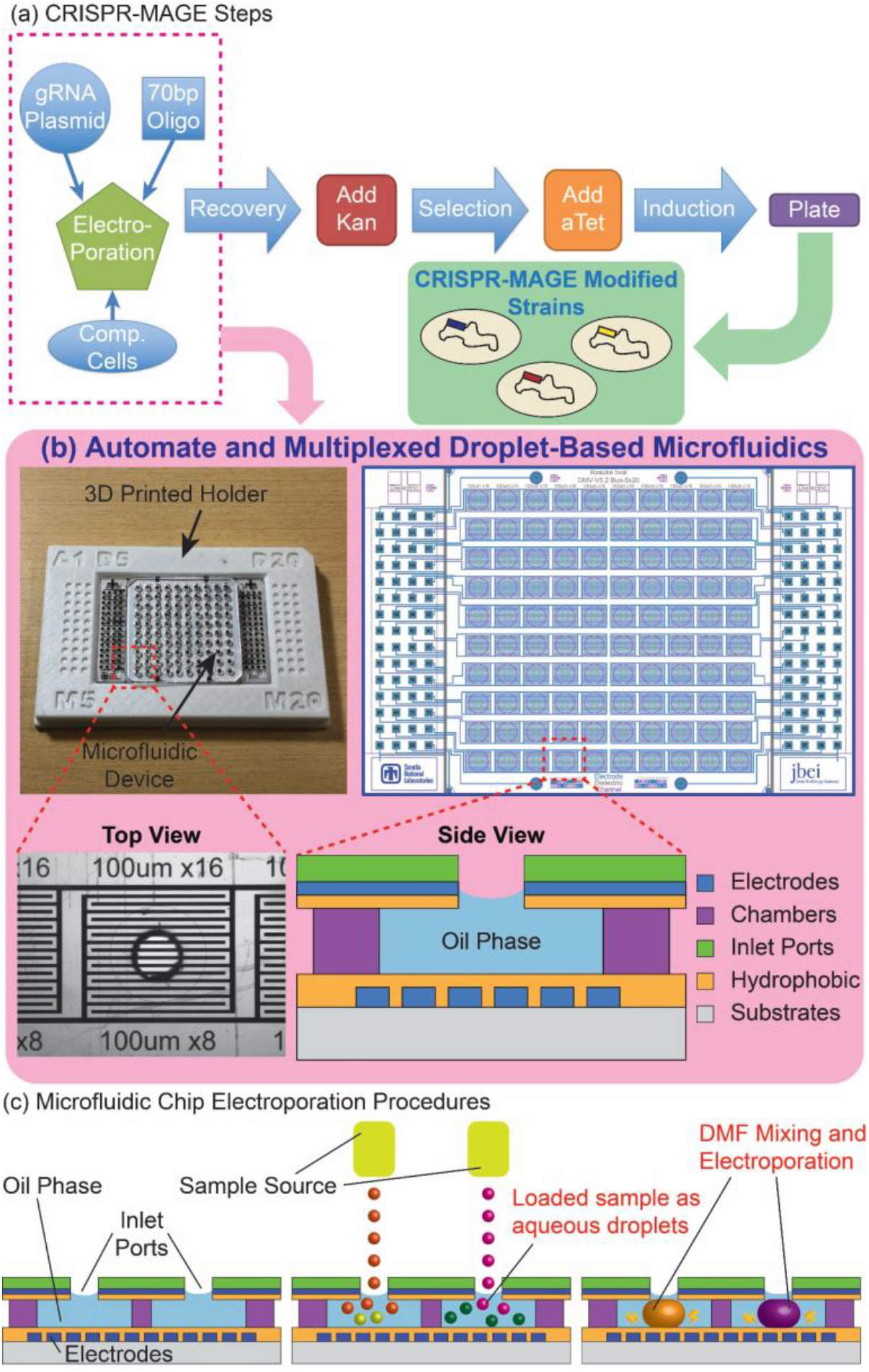
The microfluidic chip enables CRISPR-MAGE recombineering in an automated and multiplexed manner. a) Shows the steps of CRISPR-MAGE with the ones in the box performed on chip. Cells are taken out of the chip after the recovery step for induction and plating. b) The microfluidic chip is shown in a 3D printed holder (top-left), the electrode pattern (top right), top-view of an individual well, and side-view schematic of a well. The chip is designed for 100 discrete reaction chambers with individually addressable electrodes for multiplexed CRISPR-MAGE recombineering, and its 384 format design can be interfaced with lab automation equipment. c) Droplets containing plasmids and cells are directly dispensed onto each chamber through the inlet port, mixed by electrowetting manipulation, and electroporated by applying voltage pulse.

## Results and Discussion

### Microfluidic Chip Design

The microfluidic chip was designed to perform multiple functions including loading of droplets from an external liquid-handling instrument such as an acoustic printer or droplet generator into an oil layer; on-demand mixing of droplets by electrowetting; and electroporation of cells within the droplets. Reagents are introduced into the chip by dispensing droplets, are kept separate until ready to mix, mixed on-demand by merging droplets by electrowetting, and transform cells by on-chip electroporation. Additional reservoirs allow recovery incubation and screening on chip.

The first step in a high-throughput operation is the introduction of a large number of samples in parallel. We designed the microfluidic chip layout to conform to 384-well microtiter plates for a scalable approach (chips can be designed and made in 1536-well format with minimal incremental cost) for loading reagents. This loading is facilitated by the integration of the chip with a commercially-available robotic loader. We use a 3D printed holder (**Fig. 1**) to align the location of the chip connecting ports to match the jetting locations of a Labcyte Echo 550 acoustic printer. This approach, however, is general, and does not require a specific instrument: it can be used with other robotic liquid handlers, acoustic printers, droplet injectors, piezoelectric inkjet printers, or electrospray depositioners. The integration with an automated loader allows for dispensing of small (200 nm) droplets directly into the microfluidics chip. The volume of reagents injected in each well is precisely controlled by the number of droplets injected into the wells as shown in **Figure 2a**. Addition of reagents *via* droplets enables on-demand mixing--the oil contains surfactant to keep droplets separate and actuation of electrodes lowers the surface tension permitting droplets to merge for reagent mixing (**Fig. 2b-c, Supplemental Movie 1**). On-demand mixing obviates the need for manual premixing of competent cells and DNA. The oil phase also prevents evaporation, a major issue in traditional low-volume microtiter plates. While the droplet can be merged with a simple pulse of direct current (DC) voltage with duration of 10-100 msec, it is preferable to apply alternating current (AC) voltage to prevent electrolysis of the aqueous medium. We used AC frequency of 80-100 kHz with 10-100 msec duration for manipulating and merging droplets based on previous reports^10,6,7^.

**Figure 2:**
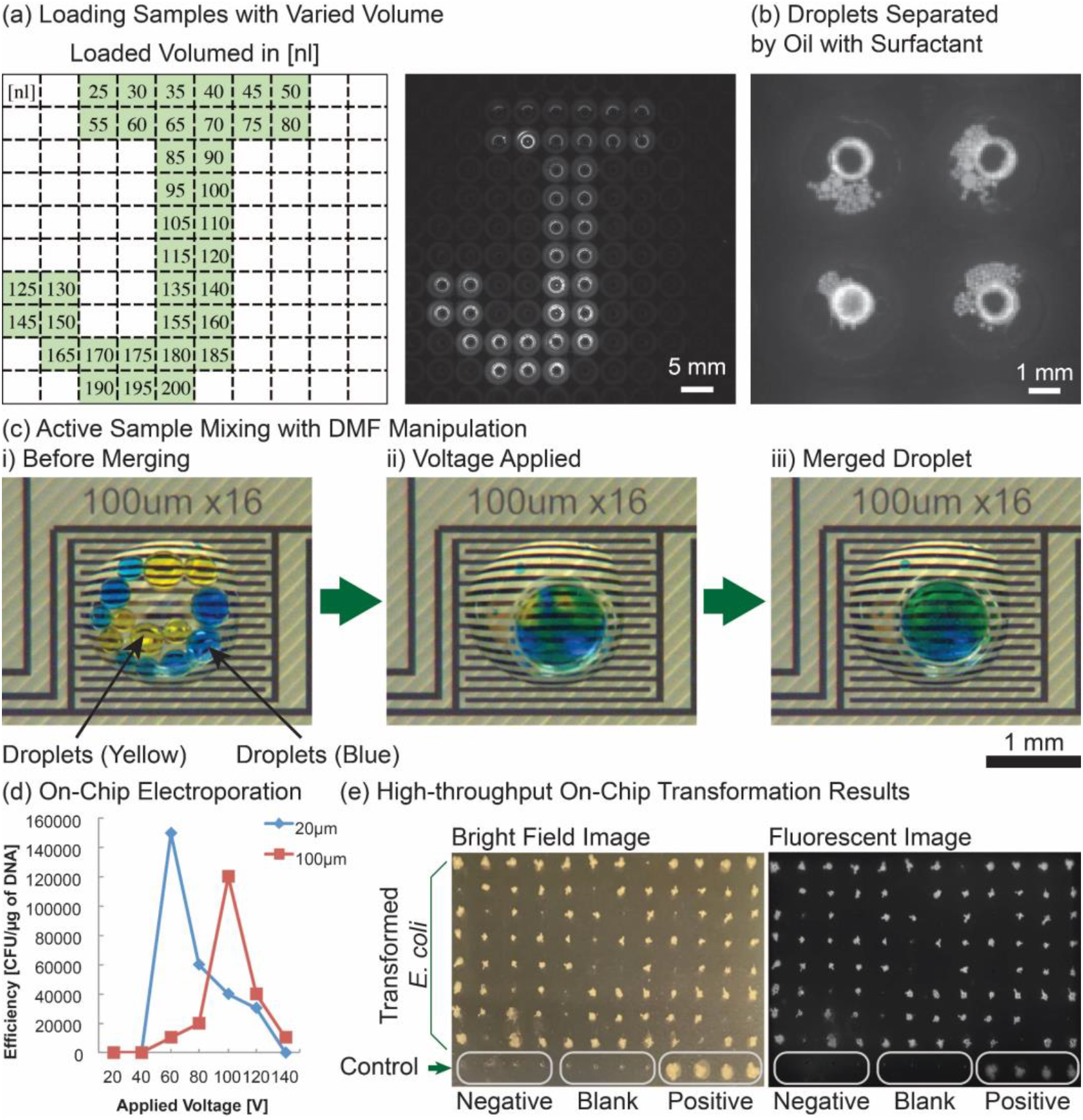
The microfluidic chip is capable of loading/mixing/electroporation at each chamber, enabling 100 discrete reactions on a single chip. (a) Sample volumes and loading sites can be programmed as desired. Green squares show where droplets were dispensed and the numbers show volume dispensed in nL. (b) Each chamber can be loaded with droplets containing different samples and the droplets are suspended in oil containing surfactant to prevent evaporation and accidental merging. (c) electrowetting enables on-demand sample mixing by merging droplets. (d) optimal voltage required for maximum transformation is dependent on the gap between electrodes. In the final chip design, we used 100 µm gap electrodes. (e) 100 parallel electroporations achieved over 80% successful transformation of E. coli with GFP plasmids. GFP was expressed by inducing with IPTG and confirmed by measuring fluorescence intensity (excitation/emission = 460nm/515-535nm).

### Electroporation in a Microfluidic Chip

Cell electroporation is potentially a more efficient process on a microfluidic chip than in cuvettes or microtiter wells on a macro-scale because the micrometer dimensions of the droplet are similar to cell diameters. Micrometer dimensions of the droplets also make diffusion-based mixing faster. We tested different electroporation conditions by fabricating microfluidic devices with two different electrode gaps, 20 µm and 100 µm, and performed transformation with different electroporation voltages for a GFP (Green Fluorescent Protein) expressing plasmid in *E. coli*. One side-effect of electroporation is electrolytically-created bubbles at the anodes (**Supplemental Figure 1, Supplemental Movies 2, 3**). The size and number of the bubbles increased with higher voltage. Shear stress from bubble formation can cause cell lysis^16^. There is an optimal point for electric field magnitude: too little and no transformation happens, too much and the cell lyses by the high field and/or lysis by bubbles. Our platform allows us to find the optimal electroporation conditions simply by changing the applied electrical pulses because each electrode is individually addressable. Electrodes with a 20 µm and 100 µm gap reached maximum efficiency at 60 V and 100 V, respectively (**Fig. 2d)**.

Our microfluidic chip configuration with individually addressable electrodes permits multiplexing of electroporation (**Supplemental Figure 2**). Multiplexing enables large numbers of DNA edits at once under identical conditions or same edits under different conditions, scaling the bioengineering process in a repeatable manner. Multiplexed electroporation, however, is difficult to perform on-chip for multiple reasons, including cross contamination, and electrical wiring and footprint limitations. Previous efforts for multiplexing have relied on serial single-plex electroporation, such as flow-through devices with a single electroporation site^10,17,18^, or the number of parallel reactions is limited to less than 10 due to physical constraints of the device configuration and electrical systems^8,19,20^. These are slow, inefficient and low-throughput processes. Our approach is inherently scalable and overcomes the cross contamination issue by physically separating each reaction site in the microfluidic chip (**Fig. 2a**). We experimentally verified our capability of multiplexed transformation by performing 100 electroporations in parallel, including 4 negative transformations, and it achieved over 80% successful transformation of *E. coli* with GFP plasmids (**Fig. 2e, Supplemental Figure 3**).

### On-Chip CRMAGE Disruption of the *galK* Gene

We implemented CRMAGE^3^ (combination of CRISPR/Cas9 with ℷ Red recombineering) in our microfluidic platform. The Cas9 protein selects against cells that do not incorporate the supplied mutated DNA oligo into their genome through ℷ Red recombineering at the position indicated by the guide RNA (gRNA). The result is a population of cells displaying edited genomes at the points determined by the oligos and gRNA. To establish proof-of-principle for on chip CRMAGE, we leveraged a classic genetic colorimetric screen. In this screen, a previously characterized point mutation to *E. coli*’s native galactokinase gene, *galK*, disables the strain from growing on a galactose medium^21–23^. When plated on MacConkey agar plates containing 1% galactose, WT strains capable of galactose fermentation turn red (no genetic change), while mutant strains are colorless (white, successful genetic change) (**Fig. 3a**). This colorimetric screen allowed us to rapidly screen the results of our engineering. CRMAGE targeting *galK* was optimized through fine tuning the concentration of the gRNA and Cas9 inducer, anhydrotetracycline (**Supplemental Figure 4**), yielding a 98 ± 3% rate of disrupting out *galK* using on-chip transformation vs 94 ± 5% in a bench process (**Fig. 3b**). As a confirmation of the colorimetric screen, *galK* mutants were also verified by Sanger sequencing (**Fig. 3c**) Our results show that CRMAGE can effectively be used on a chip to introduce point mutations, increasing the feasibility of a high-throughput CRMAGE platform.

**Figure 3:**
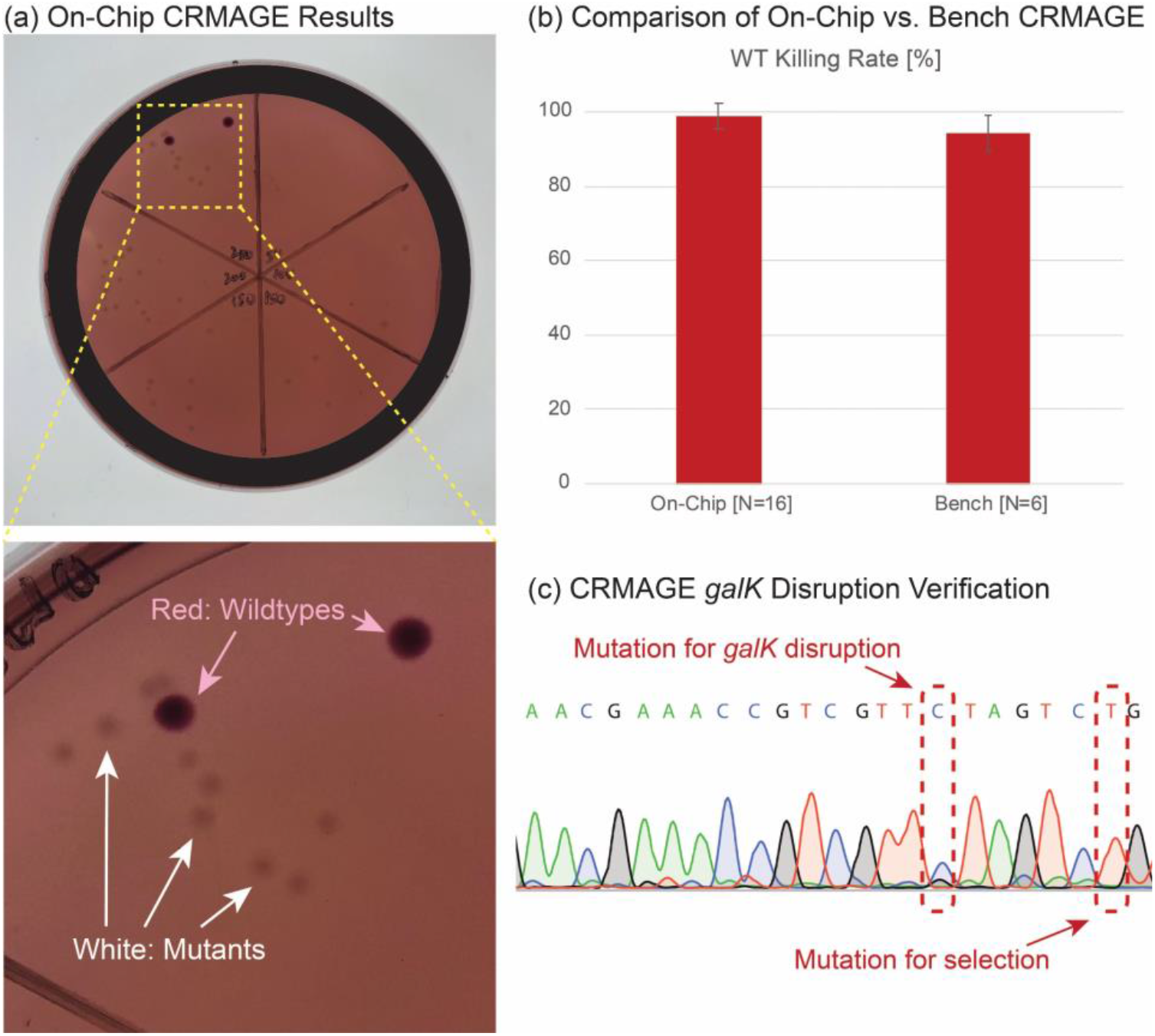
The microfluidic chip was able to use CRMAGE to disrupt galK with over 98 ± 3% success rate. (a) Mutants from on-chip electroporation are plated on MacConkey agar plates, white colonies indicating successful galK disruption, red colonies indicating wildtypes. (b) Killing rate of wildtype (WT) is similar between on-chip process and benchtop process,with both exceeding 90% success rate. Killing rates are calculated by counting the white colonies (i.e., mutants with galK disruption) over total numbers of colonies including the red colonies (i.e., wildtype). Error bars denote standard deviation of biological replicates (N=16 for on-chip process, and N=6 for benchtop process). (c) CRMAGE mutation for galk disruption is verified by Sanger sequencing. One of the mutations in the oligo produces galK disruption, while the other mutation is required for Cas9 selection since it is in the PAM region.

### On-Chip Pathway Optimization of Indigoidine-Producing Strain

To demonstrate the utility of microfluidic-CRMAGE, we applied the method to modify an *E. coli* strain producing indigoidine. To generate a stable, indigoidine producing strain, we chromosomally integrated the heterologous genes required for indigoidine production in *E. coli*: *bpsA*, the gene encoding the non-ribosomal peptide synthetase (NRPS) converting glutamine to indigoidine, and *sfp*, a 4′-phosphopantetheinyl transferase required for the activation of BpsA under control of the inducible, strong T7 promoter (**Fig. 4a**)^11,12,24^.

**Figure 4:**
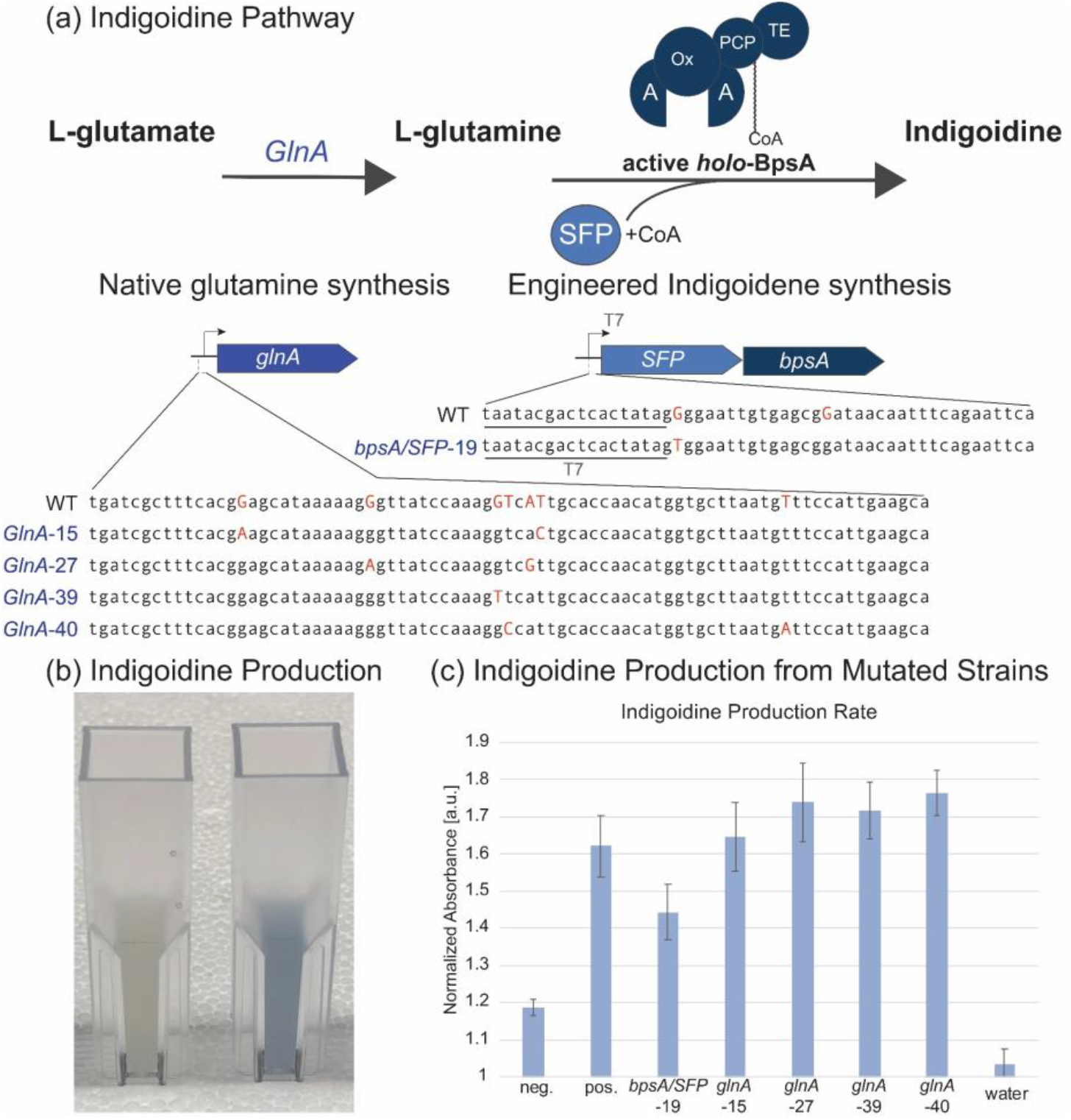
The microfluidic chip allows for automated genome modification resulting in indigoidine production changes. (a) two genetic targets were selected in the indigoidine pathway (sfp/bpsA) and four targets that affect the supply of glutamine, precursor for indigoidine production (glnA). (b) Indigoidine producing strain indicates bright blue color with highest absorbance at 615 nm. (c) The genetic modifications impact production of indigoidine (quantified by normalizing absorbance at 615 nm by 800 nm to minimize any background noise). Error bars denote standard deviation of biological triplicate.

We chose six genetic targets for point mutations (**Table 1**): two targeting the engineered pathway (*sfp* and *bpsA*) and four targeting the native glutamine pathway (*glnA*), which is the precursor for the indigoidine pathway. These sites were selected because they have a PAM site downstream and relatively few off-target sites. The results of these genomic modifications are verified by Sanger sequencing (**Supplemental Figure 5**), and can be seen reflected in the final production of indigoidine (**Fig. 4b**). The modification in the native glutamine pathway for the glnA-40 mutant produces an increase in production whereas modifications for glnA-27 and glnA-39 produce smaller increase. The bpsA/SFP-19 mutant produces a decrease in production and the glnA-15 mutant produces no change (**Fig. 4c**).

### Conclusions

We have developed a droplet-based microfluidic system capable of 100 transformations in parallel. Transformation conditions were optimized by expression of GFP in *E. coli* using electrodes with different gaps as a function of applied voltage. We adapted the CRMAGE gene-editing protocol for point mutations, and on-chip mutations disrupting *galK* achieved > 90% success rate, comparable to the results obtained with benchtop protocol. As a demonstration of our platform to optimize biosynthetic pathways, we produced 6 different mutations of indigoidine-producing *E. coli* using CRMAGE. The automated platform for multiplexed transformation holds the promise of accelerating the design-build-test-learn cycle and optimizing biosynthetic pathways. This technology could provide the technological basis for self-driving bioengineering labs^25–27^, which couples automated experimentation and AI systems that propose experiments and gauge resulting data to accelerate the bioengineering process ^28,29^.

## Materials and Methods

### Materials and Fabrication

For the fabrication process of the devices, we purchased transparent film photomasks printed at Fine Line Imaging (Colorado Springs, CO), and reagents including SU-8 5, SU-8 2075, S-1811 and SU-8 Developer from Microchem (Newton, MA), gold-coated glass substrates and chromium-coated glass substrates from Telic (Valencia, CA), indium tin oxide (ITO) coated glass slides from Delta Technologies (Stillwater, MN), MF-321 positive photoresist developer from Rohm and Haas (Marlborough, MA), CR-7 chromium etchant from OM Group (Cleveland, OH), AZ-300T photoresist stripper from AZ Electronic Materials (Somerville, NJ), Aquapel from TCP Global (San Diego, CA), and poly (dimethylsiloxane) (PDMS) from Dow Corning (Midland, MI).

Microfluidic chips are fabricated using lithography processes. First, we pattern electrodes for electroporation and electrowetting on metal-coated glass substrates using a mask aligner (OAI, Model 200). Photoresist (S-1818, Shipley) are developed with MF-321 developer after 5 seconds of UV exposure, then electrodes are etched with acids with gold etchant, CR-7 chromium etchant, or hydrochloric (HCl) acid for ITO. Electrodes yield approximately 100nm-thick after etching, and residual photoresist is stripped with an AZ stripper. We hard bake the substrates at 150 ºC for 15 minutes, then clean the surface with oxygen plasma (SERIES 800 MICRO RIE, Technics) for 2 minutes. Next, SU-8 3005 is spin coated with spinner (CEE Model 100 Spin Coating System, Brewer Science) to reach 5 µm thickness for the dielectric layer. Opening windows for electroporation are patterned with a mask aligner and developed with SU-8 developer.

For the chamber layer, we use medical grade pressure sensitive adhesive (PSA) with double-sided adhesives (Medical Tape, 3M) with roughly 100 µm in thickness. PSA layers are cut in the shape of chambers using the stencil cutter (Cameo 3, Silhouette). For the top layer, we utilize ITO-coated PET film (Indium Tin Oxide Coated PET, Sigma-Aldrich). Connecting ports with a diameter of 1 mm are milled with a computer numerical control (CNC) milling machine (Othermill, Bantam Tools). After aligning the PSA layer with the bottom substrates under a microscope, top PET film is placed onto the PSA layer. For the incubation chamber, PDMS slab is punched with a diameter of 3 mm puncher (Harris Uni-Core, Ted Pella).

For the biological samples and reagents, we purchased electro-competent *Escherichia coli* (*E. coli*) cells (MG1616) from New England Biolabs (Ipswich, MA). We obtained plasmids (GFP plasmids, pZS4Int-tetR, pMA7CR_2.0, and pMAZ-SK) for *E. coli* from public registry at the Joint BioEnergy Institute (https://public-registry.jbei.org/folders/610)^30^.

### Microfluidic Chip Design and Operation

**Figure 1** shows the design of the chips with 100 reaction sites. Microfluidic chips consist of four layers; 1) glass substrates with patterned electrodes for electroporations and DMF manipulation, 2) PSA layer for the reaction chambers, 3) top layer with ITO electrodes and connecting ports, and 4) PDMS chambers for additional incubation medium (**Supplemental Figure 6**). Reaction chambers are prefilled with oil, and droplets containing biological samples are loaded onto the chambers with external sample sources. Applied voltage to the electrowetting electrodes initiates the mixing of droplets, and electroporations are performed by applying exponential decay pulse to the pairs of electrodes in contact with the droplets. After the electroporation, droplets can be incubated either on chip by immediately introducing the recovery medium or taken to the larger volume of recovery medium in 96 plates^31^.

Liquid samples are loaded onto microfluidic chips in the form of droplets containing biological parts. Droplets with the same size scale of microfluidic channels (nanoliter∼microliter) are dispensed onto the substrates (*e*.*g*., glass, PDMS, Printed Circuit Board (PCB), etc.) of microfluidic devices either directly or remotely from the sample source (*e*.*g*., robotic liquid handler, acoustic printer, droplet injector, piezoelectric inkjet printer, electrospray deposition). Dispensing could be achieved by either directly targeting onto the microfluidic devices in the air environment, or in the oil environment through connecting ports. Sample loading process has been successfully demonstrated to load multiple aqueous samples onto a pre-sealed device *via* connecting ports without merging (**Supplemental Figure 7**). **Supplemental Figure 8** shows the fabricated microfluidic chip with a 3D printed holder to align the location of the connecting ports to match with the jetting locations.

### CRMAGE Gene-Editing

#### CRMAGE Protocol

We utilized a modified version of CRMAGE protocol developed by Ronda *et al*. ^*3*^ and its simplified process flow is shown in the subset of **Figure 1**. 100-200 ng of pMAZ-SK plasmid and 0.5 µL (10 pmoL/µL) of oligo are added to 50 µL of electrocompetent cells on ice. Electroporation pulses were applied using a commercial electroporator (Bio-Rad, Gene Pulser Xcell, and BTX, ECM-350) with a settings of voltage = 1800V (for benchtop), resistance 200 Ω, capacitance = 25 µF. Immediately after the electroporation, transformed cells are transferred to 1 mL of recovery medium of Lennox with 100 μg/mL carbenicillin (to maintain pMA7CR_2.0) and 35 μg/mL chloramphenicol (to maintain pZS4Int-tetR) and kept incubated for recovery at 37 ºC with shaking. After 2 hours of recovery incubation, kanamycin was added to reach a concentration of 50 μg/mL and the culture was incubated for additional 3 hours for selection. After 3 hours of selection incubation, anhydrotetracycline (atet) (200 ng/mL) was added and the cells were grown overnight at 37 °C shaking. Then, the cells are plated on agar plate with 50 μg/mL kanamycin, 100 μg/mL carbenicillin, and 35 μg/mL chloramphenicol and kept incubated at 37 ºC until the colonies gets visible, which typical takes approximately 12 hours.

#### CRMAGE Targeting Loci for Indigoidine Pathway Optimization

We adapted the on-chip CRMAGE process to target indigoidine pathways in *E. coli*. We engineered a wildtype strain (MG1655) to chromosomally express the genes required for indigoidine production (*bpsA* and the activator protein *sfp*) from the T7 promoter, a very strong and commonly used promoter in synthetic biology. We designed oligos and gRNA sequence for CRMAGE plasmids targeting these T7 promoters, aiming to modify gene expression of both pathway genes (*bpsA* and *sfp*) simultaneously. Additionally, we sought to investigate the impact of changing the availability of the indigoidine precursor glutamine on the production of indigoidine. Thus, we included mutations that target the native promoter driving the glutamine synthetase (*glnA*), which converts glutamate into glutamine. The designed set of oligo and gRNA sequences are shown in **Table 1**. First two sets target *sfp/bpsA*, and the other four sets target *glnA*. Following the CRMAGE guidelines from Ronda *et al*., the indigoidine-producing strain, modified MG1655, is mutated. These mutations were screened by antibiotics (kanamycin), and following selection with atet-induced Cas9 and gRNA expression. Indigoidine is produced by induction with isopropyl β-D-1-thiogalactopyranoside (IPTG). Induced cells are incubated at 30 ºC with 200 rpm for optimal production condition. Since indigoidine has strong peak absorbance at 615 nm wavelength, production rate can be quantified by measuring 615 nm absorbance with spectrophotometer as shown in **Supplemental Figure 9**. Results clearly indicate the successful production of blue pigment. The indigoidine production rate for each mutant is quantified by normalizing the 615 nm absorbance with 800 nm absorbance to minimize any background noise from unrelated molecules and to prevent pipetting errors in volume.

#### CRMAGE target selection

In order to create CRISPR gRNA, the entire JBEI-19353 strain genome was searched for PAM (NGG subsequences) on both the forward and reverse strands which are within the neighborhood of the desired genome modification site. The neighborhood in this case was chosen to be an interval of 80 bases (the size of the repair template). Upon identification of a PAM site a corresponding N20 was constructed as a 20 base region of homology with the complementing strand. These N20 sequences were then scored by doing a homology search against the rest of the genome using a fuzzy regex search allowing an off target match when the N20 deviated by fewer than 2 bases from the recognized target sequence. Six sequences were selected with the fewest off target matches found in the rest of the genome. After the design of the N20 sequences, repair templates 80 bases in length were created with the desired change integrated into them. Full code can be found in the supplementary jupyter notebook “MakeCRISPRgRNA.ipynb”.

## Supporting information

Supplementary Jupyter Notebook

## Acknowledgements

This work was part of the DOE Joint BioEnergy Institute (http://www.jbei.org) supported by the U.S. Department of Energy, Office of Science, Office of Biological and Environmental Research, through contract DE-AC02-05CH11231 between Lawrence Berkeley National Laboratory and the U.S. Department of Energy. HGM is also supported by the Basque Government through the BERC 2014-2017 program and the Spanish Ministry of Economy and Competitiveness MINECO through the BCAM Severo Ochoa excellence accreditation SEV-2013-0323. Sandia National Laboratories is a multimission laboratory managed and operated by National Technology and Engineering Solutions of Sandia, LLC., a wholly owned subsidiary of Honeywell International, Inc., for the U.S. Department of Energy’s National Nuclear Security Administration under contract DE-NA-0003525. The United States Government retains and the publisher, by accepting the article for publication, acknowledges that the United States Government retains a nonexclusive, paid-up, irrevocable, worldwide license to publish or reproduce the published form of this manuscript, or allow others to do so, for United States Government purposes.

The authors thank Tijana Radivojevic for helpful discussions and Nurgul Kaplan for the help with automated liquid handler operations.

## Author Information

### Conflicts of Interest

N.J.H declares financial interests in TeselaGen Biotechnologies and Ansa Biotechnologies.

### Contributions

K.I., M.W., M.G., L.W., D.A., N.J.H., P.D.A., A.M., H.G.M. and A.K.S. conceved the idea and planned experiments. K.I., J.S., P.W.K. and W.R.G. designed and fabricated the microfluidic chips. K.I., M.W., M.G. and D.A. developed CRMAGE protocol for indigoidine producing strain. M.W., Z.C., A.M. and H.G.M. designed the loci targets for indigoidine pathway optimization. K.I., M.W., M.G., L.W., and D.A. carried out the experiments. K.I., M.W., M.G., L.W., N.J.H., P.D.A., A.M., H.G.M., and A.K.S. wrote the manuscript.

