## Supplementary Information for "Scalable and Automated CRISPR-Based Strain Engineering Using Droplet Microfluidics"

[1]Technology Division, [2]Biofuels and Bioproducts Division, Joint BioEnergy Institute, 5855 Hollis Street, Emeryville, CA 94608, United States; [3]Sandia National Laboratories, 7011 East Avenue, Livermore, CA 94550, United States; [4]Biological Systems and Engineering Division, [5]Molecular Biophysics and Integrated Bioimaging Division, Lawrence Berkeley National Laboratory, Berkeley, CA 94720, United States; [6]Bioengineering Department, University of California, Berkeley, Berkeley, CA 94720, United States; [7]BCAM, Basque Center for Applied Mathematics, Bilbao 48009, Spain

*Co-corresponding authors

**Table 1**: Designed oligo and gRNA targeting indigoidine pathway

| Mutation # | Target | N20 | Repair Template |
|---|---|---|---|
| #1 | *SFP/bpsA* | AAATTAATACGACTCACTAT | ATCGATCTCGATCCCGCGAAATTAATACGACTCACTATAGAGGAATTGTGAGCGGATAACAATTTCAGAATTCAAAAGA |
| #2 | *SFP/bpsA* | CACTATAGGGGAATTGTGAG | ATCGATCTCGATCCCGCGAAATTAATACGACTCACTATAGAGGAATTGTGAGCGAATAACAATTTCAGAATTCAAAAGA |
| #3 | *glnA* | TCCCTTTGTGATCGCTTTCA | TGATCGCTTTCACGAAGCATAAAAAGGGTTATCCAAAGGTCACTGCACCAACATGTGCTTAATGTTTCCATTGAAGCA |
| #4 | *glnA* | CGCTTTCACGGAGCATAAAA | TGATCGCTTTCACGGAGCATAAAAAGAGTTATCCAAAGGTCGTTGCACCAACATGGTGCTTAATGTTTCCATTGAAGCA |
| #5 | *glnA* | GCATAAAAAGGGTTATCCAA | TGATCGCTTTCACGGAGCATAAAAAGGGTTATCCAAAGTTCATTGCACCAACATGGTGCTTAATGTTTCCATTGAAGCA |
| #6 | *glnA* | CAAAGGTCATTGCACCAACA | TGATCGCTTTCACGGAGCATAAAAAGGGTTATCCAAAGGCCATTGCACCAACATGATGCTTAATGTTTCCATTGAAGCA |

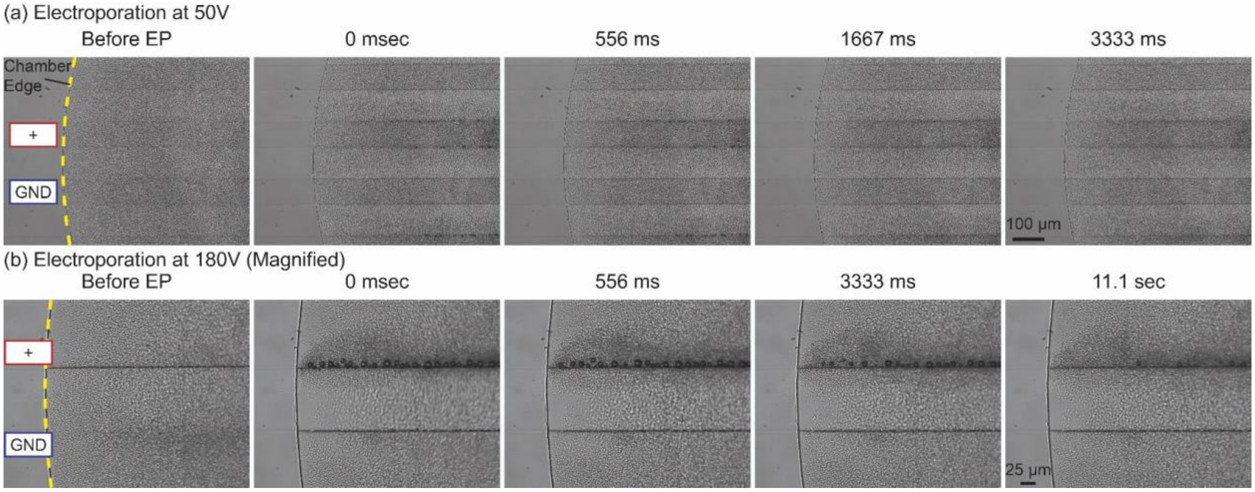


***SI Figure 1****: Larger bubbles are formed at the edges of the positive electrodes when higher voltage is applied. (a) Small bubbles are formed at 50 V with 100 µm electrode gap and they disappear approximately after 3 seconds. Average diameter of the bubbles were 4.7 µm ± 1.1 µm (N = 20) (b) More of larger bubbles are formed at 180 V with 100 µm electrode gap and it takes more than 11 seconds to disappear. Average diameter of the bubbles were 8.4 µm ± 1.6 µm (N = 23) Yellow dashed line indicates the edge of the reaction chambers and electrocompetent cells are initially suspended uniformly inside these chambers. Horizontal lines are the boundary of the transparent (ITO) electrodes (square boxes outlined in blue indicate the ground electrodes and square boxes outlined in red indicate the high-voltage electrodes).*

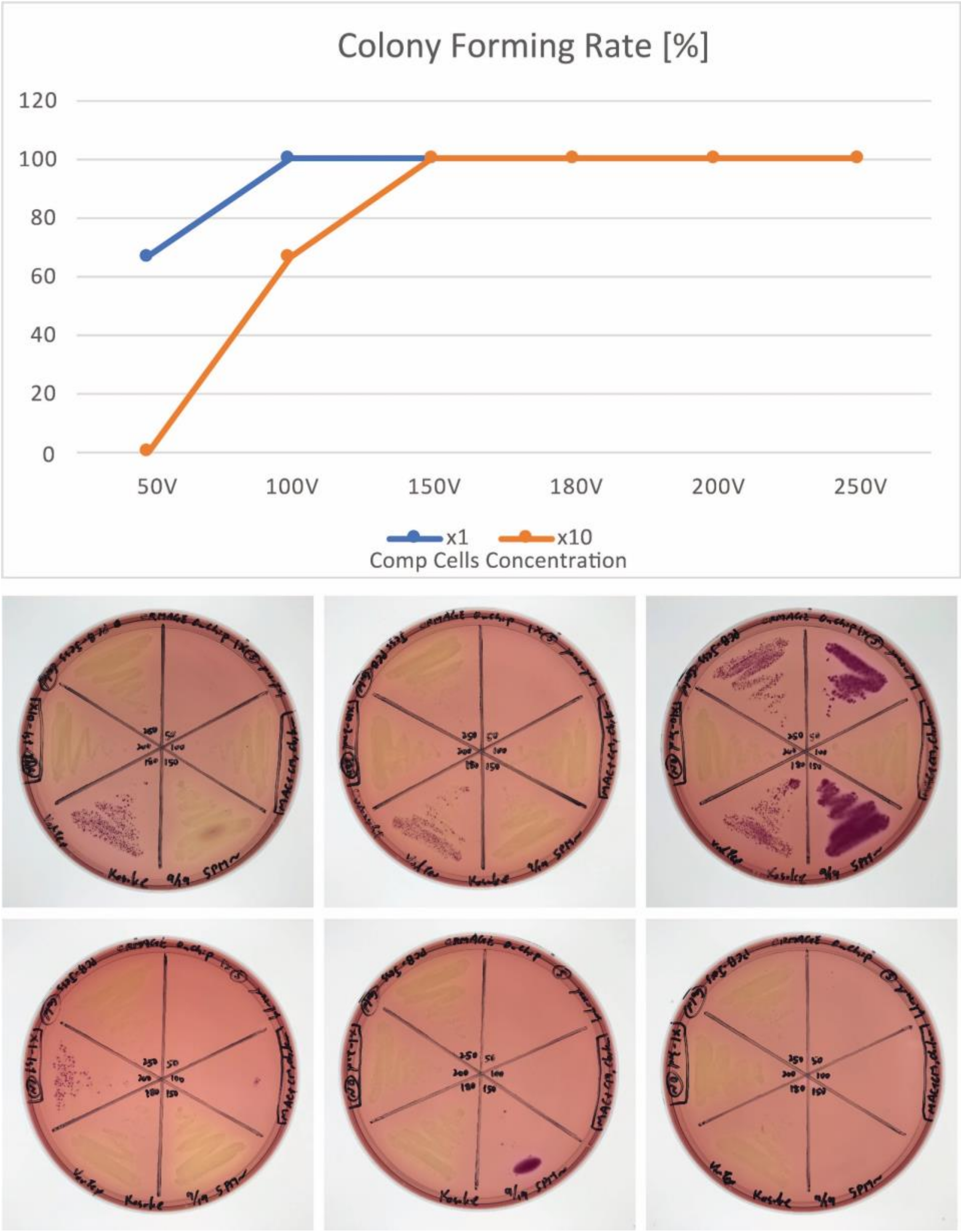


***SI Figure 2****: Our microfluidic chip enables multiplexed experiments on a single chip. Multiplexed experiments are successfully demonstrated for* galK *disruption with different electroporation conditions. We tested six different conditions of electroporation voltages (50, 100, 150, 180, 200, 250 [V]) with two different concentrations of the competent cells (regular concentration: x1, and 10 times denser cells: x10) with triplicates. Results indicate successful colony forming per each experiment.*

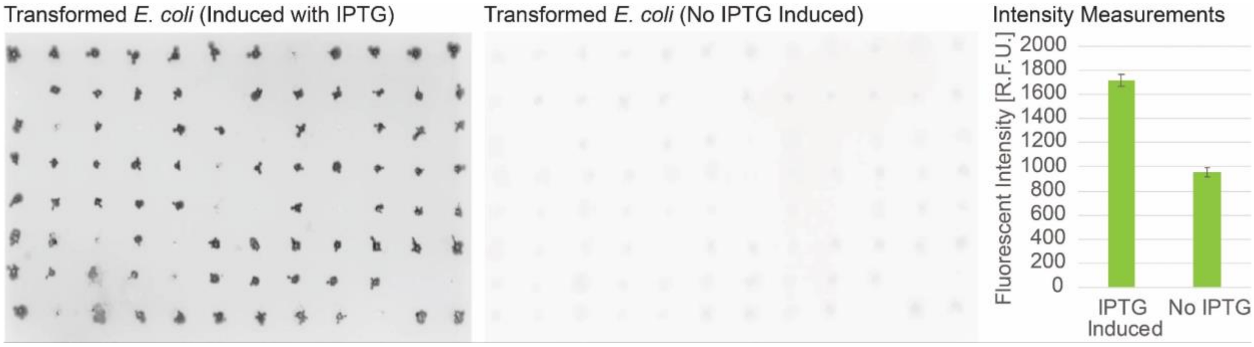


***SI Figure 3****: Fluorescent images of transformed E. coli with GFP plasmids for fluorescent intensity analysis. Left image is E. coli colonies induced with IPTG, showing significantly high fluorescent intensity. Right image is E. coli colonies without IPTG added, showing low fluorescent intensity, with little background due to the leaking of the promoter to allow GFP to express in small amounts without IPTG. Error bars denote standard deviation of biological replicates (N=96)*

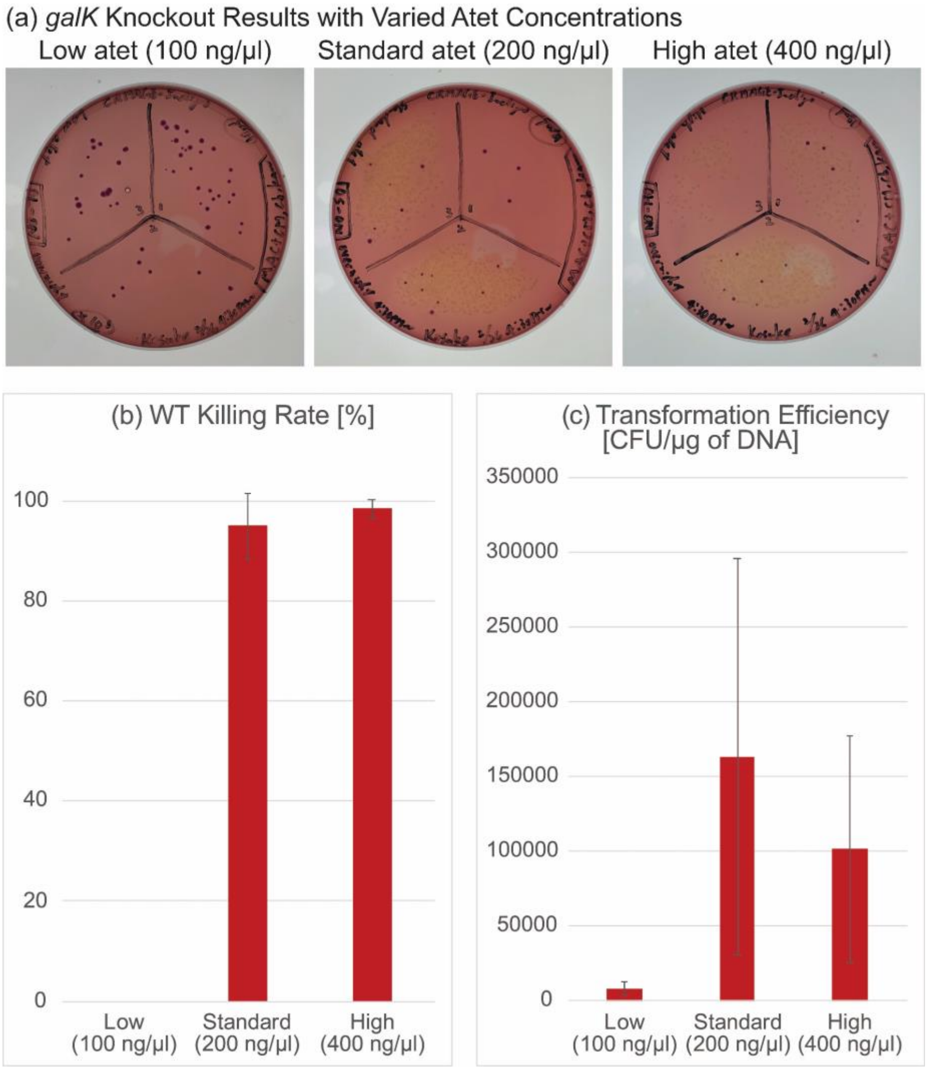


***SI Figure 4****: Anhydrotetracycline (atet) inducer concentration is optimized to get maximum wildtype killing rate and overall transformation efficiency. (a) Results of CRMAGE* galK *disruption using different atet concentrations: low (100 ng/µL), standard (200 ng/µL), and high (400 ng/µL) (b) Both of standard and high concentration achieved high wildtype killing rate, while low concentration did not work to kill the wildtypes. (c) Standard concentration achieved highest overall transformation efficiency.*

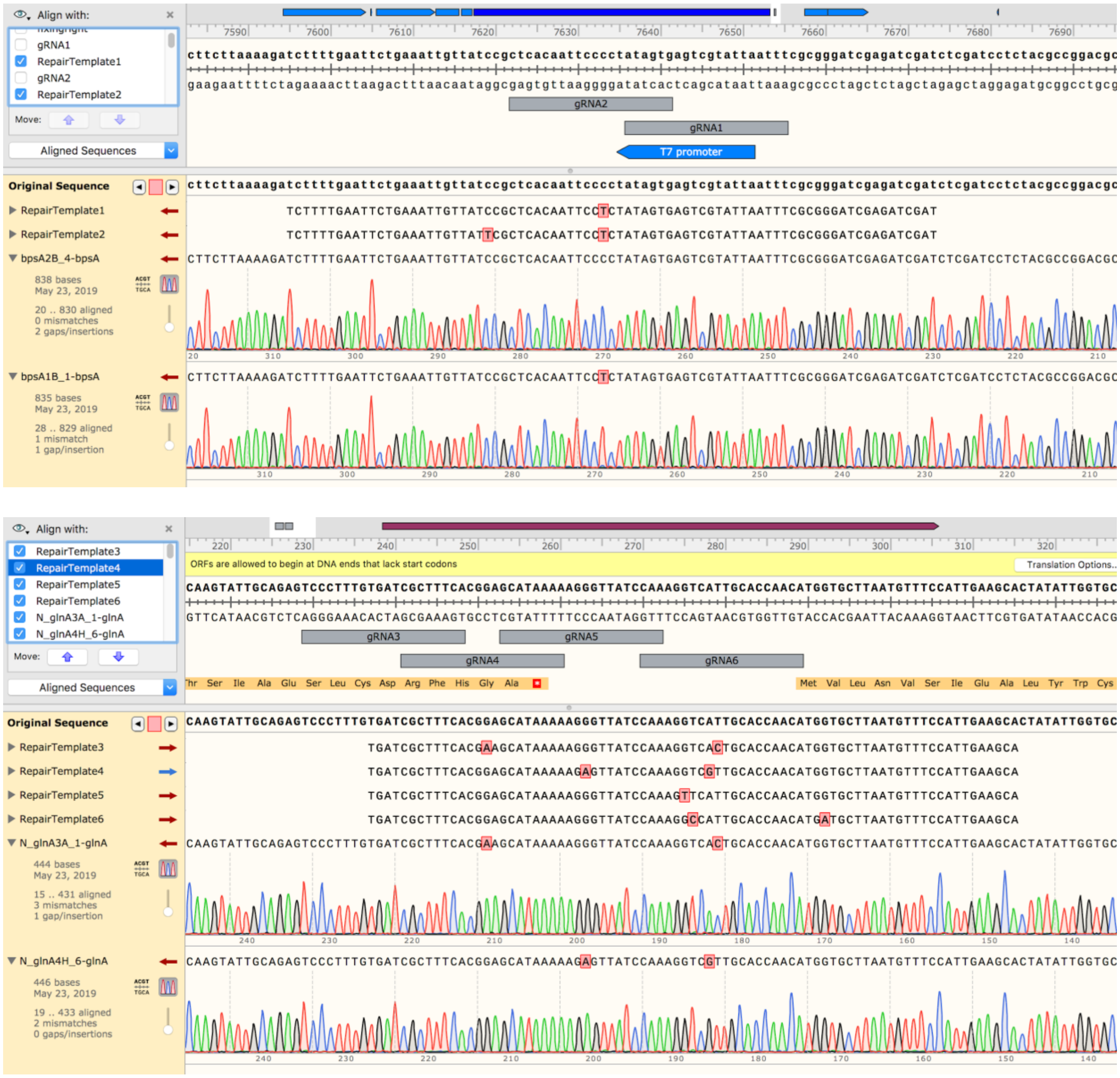


***SI Figure 5****: The CRMAGE point mutations targeting the indigoidine pathway were verified by Sanger sequencing.*

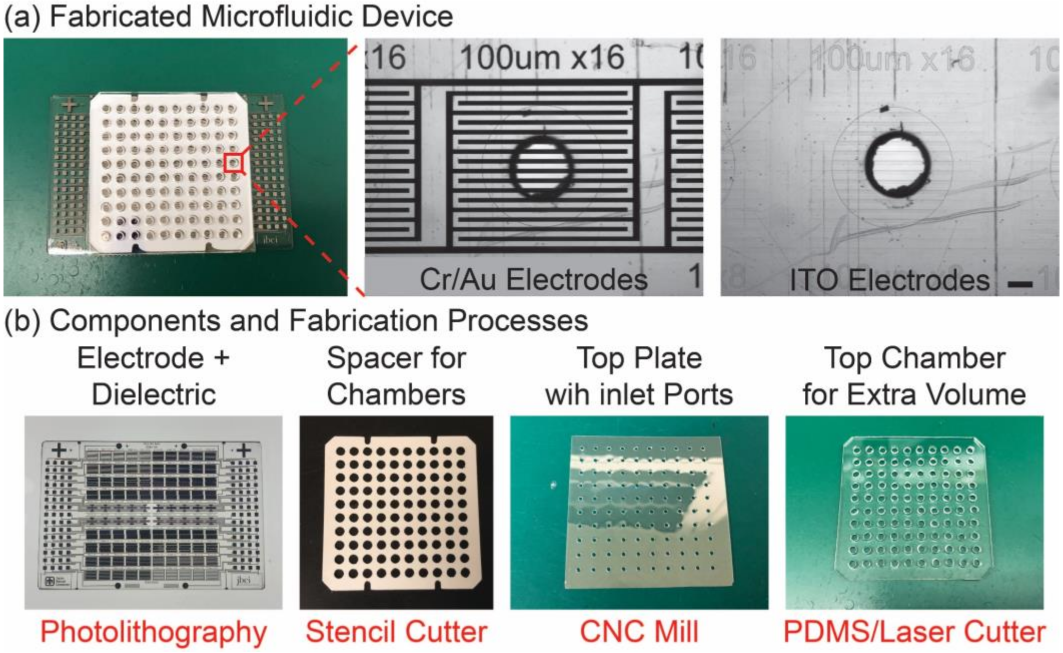


***SI Figure 6**: **The microfluidic chip can be fabricated by standard prototyping procedures**. Here, we show fabricated microfluidic devices with 100 reaction chambers in 384 well format, and components with fabrication processes. (a) The electrode material can be either standard Chrome/Gold electrodes or Indium Tin Oxide (ITO) for optical transparency for compatibility with microplate readers. Scale bars = 500 µm. (b) The microfluidic chip consists of four layers and each layer can be made with standard prototyping methods. First layer is electrowetting/electroporation electrodes patterned with photolithography. Second layer is a 100 µm-thick chamber layer that is cut* via *a stencil cutter. Top plate is a PET film with access inlet ports drilled with CNC mills. The top additional chambers can be made with punched PDMS or laser cut polymers to provide additional volume.*

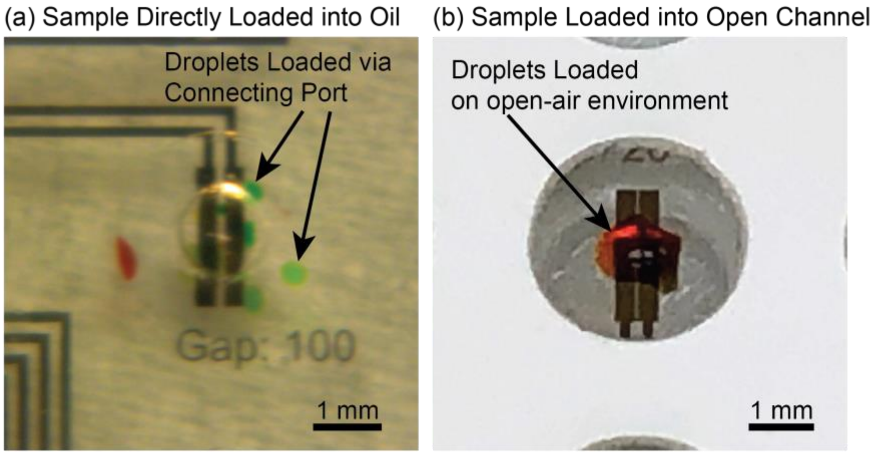


***SI Figure 7:*** *Demonstration of aqueous sample loading with an acoustic printer. (a) Droplets are separated when the channel is pre-filled with an oil phase with surfactant. (b) When the channel is open without oil, droplets are merged immediately after loading, and they evaporate quickly due to the high surface-volume ratio.*

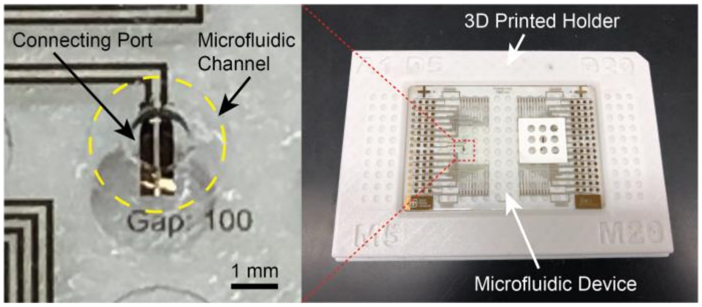


***SI Figure 8:*** *Fabricated microfluidic device with connecting ports. Yellow dashed circle shows the microfluidic channel.*

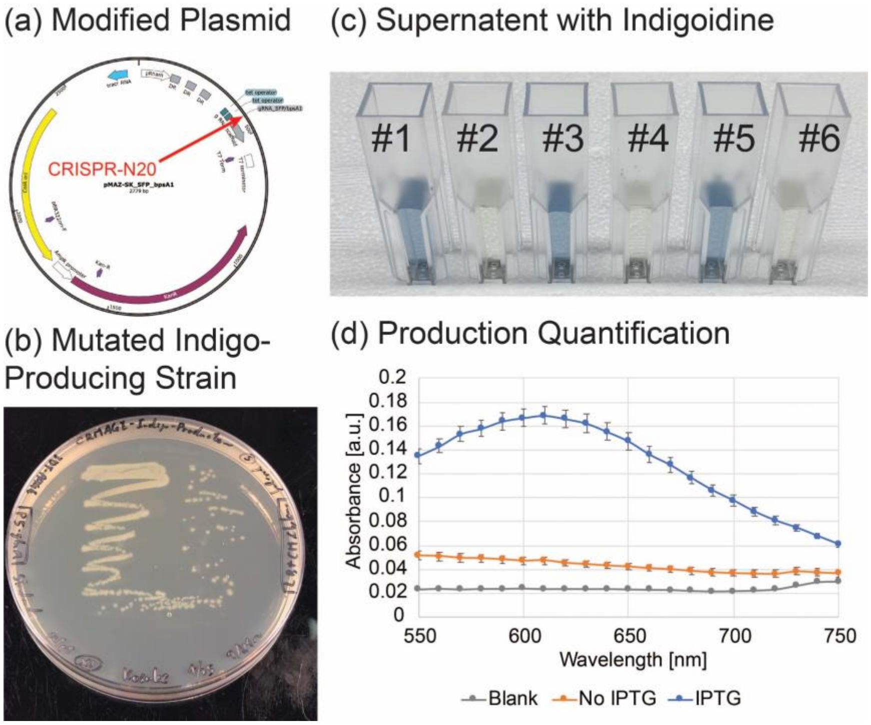


***SI Figure 9**: Benchtop indigoidine production results for quantifying the production rate by absorbance. (a) Modified pMAZ-SK plasmid with gRNA sequence to target indigoidine pathway. (b) Mutated indigo-producing strain. (c) Indigoidine-producing strain is centrifuged to separate indigoidine containing supernatant. (Samples #1, 3, 5 are from indigoidine production, and samples #2, 4, 6 are without indigoidine production) (d) Successful indigoidine production quantified by absorbance. Produced indigoidine shows highest absorbance at 615 nm. Error bars denote standard deviation of biological triplicate.*

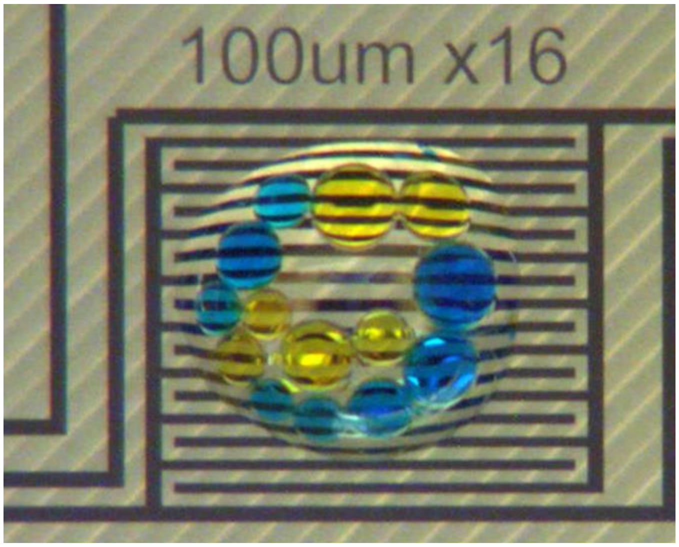


***SI Movie 1**: Yellow colored droplets and blue colored droplets are mixed by applying voltage.*

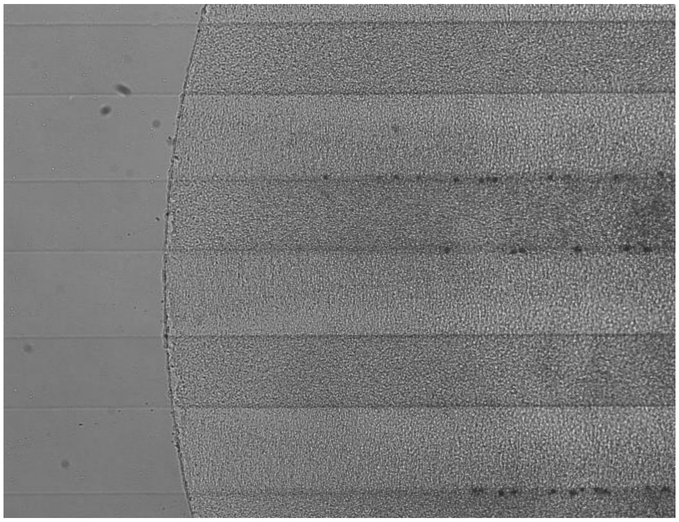

***SI Movie 2**: Small bubbles are formed at 50 V with 100 μm electrode gap and they disappear approximately after 3 seconds*

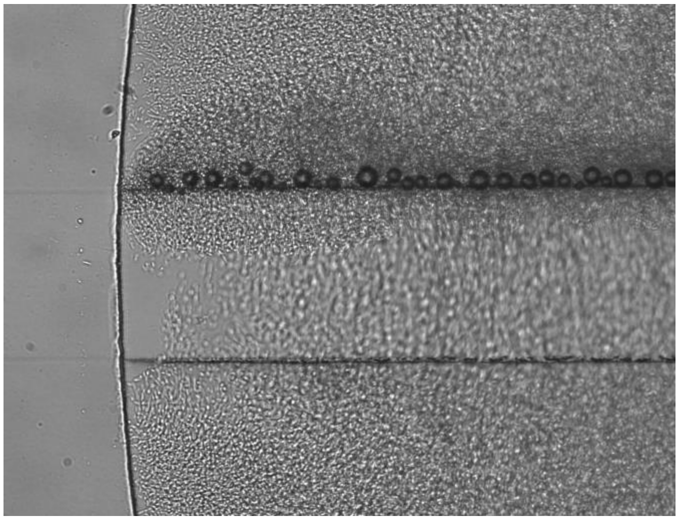

***SI Movie 3****: larger bubbles are formed at 180 V with 100 µm electrode gap and it takes more than 11 seconds to disappear*
