## Supplementary Jupyter Notebook for "Scalable and Automated CRISPR-Based Strain Engineering Using Droplet Microfluidics"

### Create CRISPR gRNA with Minimal Off Target Effects

In this notebook we take a genome, a list of target modification pairs, and oligo length and return a set of crispr guide RNAs, their associated DNA sequence, The Repair Template, and an off target score.

In [1]:
```python
from Bio import SeqIO
from Bio.Seq import Seq
from Bio.Alphabet import DNAAlphabet
from Bio import pairwise2
from itertools import chain
import math
import re
import regex

import numpy as np
import pandas as pd
from IPython.display import display

import matplotlib.pyplot as plt
```

In [2]:
```python
#Read in Genome
for record in SeqIO.parse("data/jbei-19353_genome.fasta", "fasta",alphabet=DNAAlphabet()):
    jbei_19353 = record.seq
```

In [3]:
```python
def mage_design(genome,in_seq,out_seq,max_template_length=80,pam_sequence=Seq('NGG',DNAAlphabet())):
    '''Looks for in_seq in the genome then designs a set of gRNA
    and repair_template to make the change along with scores.
    '''

    pattern = re.compile('.GG')

    #Find In_Seq (Try Forward and Reverse)
    seq_start = genome.find(in_seq)

    #Find Differences between In_Seq and Out Seq
    alignments = pairwise2.align.localxx(in_seq,out_seq)
    in_align = alignments[0][0]
    out_align = alignments[0][1]

    n = len(in_align)
    f_dash = False
    b_dash = False
    for f in range(n):
        b = n - f - 1
        if ((in_align[f] == '-') | (out_align[f] == '-')) & ~f_dash:
            left_ha_end = f - 1
            f_dash = True

        if ((in_align[b] == '-') | (out_align[b] == '-')) & ~b_dash:
            right_ha_start = b + 1
            b_dash = True

    repair_core = out_align[(left_ha_end + 1):(right_ha_start)].replace('-','')
    right_ha_start = right_ha_start - len(repair_core)

    ha_len = math.floor((max_template_length - len(repair_core))/2)
    left_ha = genome[(seq_start + left_ha_end - ha_len + 1):(seq_start + left_ha_end+1)]
    right_ha = genome[(seq_start + right_ha_start):(seq_start + right_ha_start + ha_len)]

    repair_template = left_ha + repair_core + right_ha
    repair_template_start = seq_start + left_ha_end - ha_len + 1
    repair_template_end = seq_start + right_ha_start + ha_len
    genome_repair_region = genome[repair_template_start:repair_template_end]

    #Find Pam Sites In Genome That can be mutated by the repair template
    match = re.finditer(pattern, str(genome_repair_region))
    pam_locations = [(m.start(0)+repair_template_start, m.end(0)+repair_template_start) for m in re.finditer(pattern,
str(genome_repair_region))]
    pam_sites = [genome[i:j] for i,j in pam_locations]

    #print(repair_template)
    #print(len(repair_template))
    #print(pam_locations)
    #print(pam_sites)

    #For Each Pam Site Create gRNA, repair_template, and Score
    designs = []
    for pam_site in pam_locations:
        #mutate pam site of repair template
        mutated_repair_template = repair_template

        #Create CRISPR Guide to Target Pam Site
        gRNA = create_gRNA(genome,pam_site)

        #Append Found GRNA
        designs.append((gRNA,mutated_repair_template,repair_template_start))

    return designs


def create_gRNA(genome,pam_site):
    '''create n20 guide RNA to target a particular pam site in the genome'''
    binding_region = genome[pam_site[0]-20:pam_site[0]]
    gRNA = binding_region.transcribe()
    return gRNA

def score_gRNA(genome,gRNA):
    '''look to see if there are any spots where the gRNA matches'''

    r = regex.compile('({}){{e<=2}}'.format(str(gRNA.back_transcribe())))

    #Perform Forward Pass
    matches = regex.finditer(r,str(genome))

    #Perform Reverse Pass
    matches = chain(matches,regex.finditer(r,str(genome.reverse_complement())))

    n = 0
    for match in matches:
        n+= 1

    return n


def print_design(genome_region,gRNA,repair_template,forward=True,ha_length=18):
    '''Prints out Repair Template To Understand the Design Visually'''
    gRNA_align = pairwise2.align.localxs(genome_region,gRNA.back_transcribe(),-10,-10)

    #Create a Unified Alignment from two independent alignments
    left_ha = repair_template[:ha_length]
    right_ha = repair_template[-ha_length:]
    left_align = pairwise2.align.localxs(genome_region,left_ha,-10,-10)
    right_align = pairwise2.align.localxs(genome_region,right_ha,-10,-10)

    repair_align = ''
    before_lh = True
    before_rh = True
    for l,r in zip(left_align[0][1],right_align[0][1]):
        if (l == '-') and (r == '-'):
            if (~before_lh) & before_rh:
                repair_align += '-'
            else:
                repair_align += ' '
        elif (l == '-'):
            repair_align += r
            before_rh = False

        elif (r == '-'):
            repair_align +=l
            before_lh = False


    #Print DNA
    if forward:
        print('   {}:back translated gRNA'.format(gRNA_align[0][1]).replace('-',' '))
        print('   {}:repair template homology'.format(repair_align))
        print('5\':{}:3\''.format(genome_region))
        print('3\':{}:5\''.format(genome_region.complement()))
    else:
        print('3\':{}:5\''.format(genome_region.complement()))
        print('5\':{}:3\''.format(genome_region))
        print('   {}:repair template homology'.format(repair_align))
        print('   {}:back translated gRNA'.format(gRNA_align[0][1]).replace('-',' '))

    print('')
    print('Repair Template:')
    print('5\':{}:3\''.format(repair_template))

    #Get Local Region
    pass
```

In [4]:
```python
#Quick Test Case
genome = Seq('TTTTAGGGCCCTAATTAGTCAGGCCATATTTTTTTGGAGGAAATCCCCCCCTAAAATTTTTTTTTTTTTTTTTTT',DNAAlphabet())
in_seq = Seq('AAATCCCCCCCTAA',DNAAlphabet())
out_seq = Seq('AAATGGGGTAA',DNAAlphabet())

#Forward Designs
designs = mage_design(genome,in_seq,out_seq,max_template_length=40)
for i,design in enumerate(designs):
    print('Forward Design {}'.format(i+1))
    print('score (lower is better): {}'.format(score_gRNA(genome,design[0])))
    print_design(genome,design[0],designs[0][1])
    print('')

#Reverse Designs
designs = mage_design(genome.reverse_complement(),in_seq.reverse_complement(),out_seq.reverse_complement(),max_templat
e_length=40)
for i,design in enumerate(designs):
    print('Forward Design {}'.format(i+1))
    print('score (lower is better): {}'.format(score_gRNA(genome,design[0])))
    print_design(genome.reverse_complement(),design[0],designs[0][1],forward=False)
    print('')
```

```
Forward Design 1
score (lower is better): 1
                 TTAGTCAGGCCATATTTTTT                                        :back translated gRNA
                             TATTTTTTTGGAGGAAAT-------TAAAATTTTTTTTTTTTTT     :repair template homology
5':TTTTAGGGCCCTAATTAGTCAGGCCATATTTTTTTGGAGGAAATCCCCCCCTAAAATTTTTTTTTTTTTTTTTTT:3'
3':AAAATCCCGGGATTAATCAGTCCGGTATAAAAAAACCTCCTTTAGGGGGGGATTTTAAAAAAAAAAAAAAAAAAA:5'

Repair Template:
5':TATTTTTTTGGAGGAAATGGGGTAAAATTTTTTTTTTTTTT:3'

Forward Design 2
score (lower is better): 1
                    GTCAGGCCATATTTTTTTGG                                     :back translated gRNA
                             TATTTTTTTGGAGGAAAT-------TAAAATTTTTTTTTTTTTT     :repair template homology
5':TTTTAGGGCCCTAATTAGTCAGGCCATATTTTTTTGGAGGAAATCCCCCCCTAAAATTTTTTTTTTTTTTTTTTT:3'
3':AAAATCCCGGGATTAATCAGTCCGGTATAAAAAAACCTCCTTTAGGGGGGGATTTTAAAAAAAAAAAAAAAAAAA:5'

Repair Template:
5':TATTTTTTTGGAGGAAATGGGGTAAAATTTTTTTTTTTTTT:3'

Forward Design 1
score (lower is better): 1
3':TTTTTTTTTTTTTTTTTTTAAAATCCCCCCCTAAAGGAGGTTTTTTTATACCGGACTGATTAATCCCGGGATTTT:5'
5':AAAAAAAAAAAAAAAAAAATTTTAGGGGGGGATTTCCTCCAAAAAAATATGGCCTGACTAATTAGGGCCCTAAAA:3'
        AAAAAAAAAAAAAATTTTA-------ATTTCCTCCAAAAAAATA                         :repair template homology
     AAAAAAAAAAAAAAAAAATTTT                                                  :back translated gRNA

Repair Template:
5':AAAAAAAAAAAAAATTTTACCCCATTTCCTCCAAAAAAATA:3'

Forward Design 2
score (lower is better): 1
3':TTTTTTTTTTTTTTTTTTTAAAATCCCCCCCTAAAGGAGGTTTTTTTATACCGGACTGATTAATCCCGGGATTTT:5'
5':AAAAAAAAAAAAAAAAAAATTTTAGGGGGGGATTTCCTCCAAAAAAATATGGCCTGACTAATTAGGGCCCTAAAA:3'
        AAAAAAAAAAAAAATTTTA-------ATTTCCTCCAAAAAAATA                         :repair template homology
        AAAAAAAAAAAAAATTTTAGG                                                :back translated gRNA

Repair Template:
5':AAAAAAAAAAAAAATTTTACCCCATTTCCTCCAAAAAAATA:3'
```

#### Finding the Correct Promoter To Use

In [5]:
```python
def levenshtein(seq1, seq2):
    size_x = len(seq1) + 1
    size_y = len(seq2) + 1
    matrix = np.zeros ((size_x, size_y))
    for x in range(size_x):
        matrix [x, 0] = x
    for y in range(size_y):
        matrix [0, y] = y

    for x in range(1, size_x):
        for y in range(1, size_y):
            if seq1[x-1] == seq2[y-1]:
                matrix [x,y] = min(
                    matrix[x-1, y] + 1,
                    matrix[x-1, y-1],
                    matrix[x, y-1] + 1
                )
            else:
                matrix [x,y] = min(
                    matrix[x-1,y] + 1,
                    matrix[x-1,y-1] + 1,
                    matrix[x,y-1] + 1
                )
    return (matrix[size_x - 1, size_y - 1])
```

In [6]:
```python
#Read in in_seq, out_seq pairs & Generate gRNA and repair templates

#SFP (Forward Strand) T7 Promoter Sequence: taatacgactcactatagg
#SFP (Forward Strand) Extended Sequence (for homology): atcgatctcgatcccgcgaaattaatacgactcactataggggaat


#bpsA (Reverse Strand) T7 Promoter Sequence: taatacgactcactatagg
#bpsA (Reverse Strand) Extended Sequence: aggatcgagatcgatctcgatcccgcgaaattaatacgactcactataggggaattgtgagcgga

#Need to Find Alternative T7 Promoter Sequences...
```

In [7]:
```python
#Create Knockdown Targets for T7 Promoters Driving Heterologous Genes!
t7_wt_promoter = 'TAATACGACTCACTATAGG'

t7df = pd.read_csv('data/T7Variants.csv')
t7df['log2 strength'] = np.log2(t7df['RNA read/DNA read'])
t7df_mu = t7df.groupby('Randomized T7 promoter sequence').mean()
t7df_mu['strength'] = t7df_mu['RNA read/DNA read']
t7df_sigma = t7df.groupby('Randomized T7 promoter sequence').std()
t7df_mu['strength std'] = t7df_sigma['RNA read/DNA read']
t7df = t7df_mu
```

In [8]:
```python
#calculate edit distance
edit_distance = lambda x: levenshtein(t7_wt_promoter,x)
t7df['edit_distance'] = t7df.index.map(edit_distance)
```

In [9]:
```python
t7df.plot.scatter(x='log2 strength', y='edit_distance', marker='.')
plt.show()
```

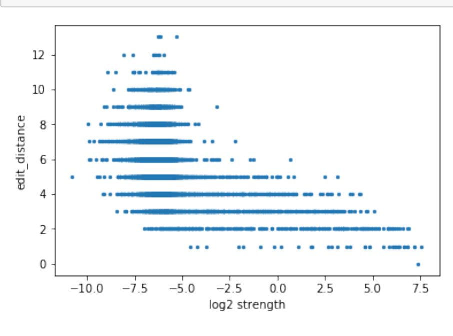


In [10]:
```python
#Find Default Sequence
wt_strength = t7df.loc[t7_wt_promoter]['log2 strength']
print(wt_strength)
t7df_knockdown = t7df.loc[(t7df['log2 strength'] > wt_strength - 2)&(t7df['log2 strength'] < wt_strength - 1)&(t7df['e
dit_distance']==1)]
display(t7df_knockdown.sort_values('RNA read/DNA read',ascending=False))
```

```
7.395045650405885
```

| Randomized T7 promoter sequence | DNA read number | RNA (barcode) read number | RNA read/DNA read | Duplication number | log2 strength | strength | strength std | edit_distance |
|---|---|---|---|---|---|---|---|---|
| TAATCCGACTCACTATAGG | 1293.000000 | 106852.5 | 82.090128 | 1.0 | 6.353379 | 82.090128 | 10.350947 | 1.0 |
| TAATACGACTCACTACAGG | 1090.000000 | 87713.0 | 80.470642 | 1.0 | 6.330391 | 80.470642 | NaN | 1.0 |
| TAATACGACTCACTATAGA | 2180.500000 | 161725.5 | 74.276858 | 1.0 | 6.214226 | 74.276858 | 3.065506 | 1.0 |
| TAATACGACTCACTATTGG | 2439.000000 | 178696.0 | 73.266093 | 1.0 | 6.195074 | 73.266093 | NaN | 1.0 |
| TAATACGACTCACTCTAGG | 850.000000 | 60535.5 | 68.043657 | 1.0 | 6.053693 | 68.043657 | 20.853013 | 1.0 |
| TAATACGACTCACTATAAG | 1375.000000 | 93507.0 | 68.005091 | 1.0 | 6.087571 | 68.005091 | NaN | 1.0 |
| TAATACGACTCACTAGAGG | 854.333333 | 50916.0 | 58.541166 | 1.0 | 5.847262 | 58.541166 | 12.489596 | 1.0 |
| TAATACGACTCGCTATAGG | 1020.500000 | 47408.5 | 47.685101 | 1.0 | 5.574770 | 47.685101 | 2.095250 | 1.0 |

In [11]:
```python
#Create Constructs
#Promoter Knockdown Chosen: TAATACGACTCACTATAGA (Should reduce expresion by about half)
wt_promoter = 'TAATACGACTCACTATAGG'
kd_promoter = 'TAATACGACTCACTATAGA'
in_seq = Seq('AAATCCCCCCCTAA',DNAAlphabet())
out_seq = Seq('AAATGGGGTAA',DNAAlphabet())

constructs = ['SFP','bpsA']
in_seqs = [Seq('atcgatctcgatcccgcgaaat{}ggaat'.format(wt_promoter).upper(),DNAAlphabet()),
           Seq('aggatcgagatcgatctcgatcccgcgaaat{}ggaattgtgagcgga'.format(wt_promoter).upper(),DNAAlphabet())]
out_seqs = [Seq('atcgatctcgatcccgcgaaat{}ggaat'.format(kd_promoter).upper(),DNAAlphabet()),
           Seq('aggatcgagatcgatctcgatcccgcgaaat{}ggaattgtgagcgga'.format(kd_promoter).upper(),DNAAlphabet())]
```

In [12]:
```python
#Create Designs for Each Construct
for construct,in_seq,out_seq in zip(constructs,in_seqs,out_seqs):
    print(construct)
    designs = mage_design(jbei_19353,in_seq,out_seq,max_template_length=80)
    for i,design in enumerate(designs):
        genome = jbei_19353[(design[-1]-30):(design[-1]+100)]
        #print(design)
        print(design[-1])
        print('Forward Design {}'.format(i+1))
        print('score (lower is better): {}'.format(score_gRNA(jbei_19353,design[0])))
        print_design(genome,design[0],designs[0][1],ha_length=37)
        print('')

    #Reverse Designs
    designs = mage_design(jbei_19353.reverse_complement(),in_seq.reverse_complement(),out_seq.reverse_complement(),max
_template_length=80)
    for i,design in enumerate(designs):
        #print(design)
        print(design[-1])
        print('Reverse Design {}'.format(i+1))
        print('score (lower is better): {}'.format(score_gRNA(jbei_19353,design[0])))
        print_design(genome.reverse_complement(),design[0],designs[0][1],forward=False,ha_length=37)
        print('')
```

```
SFP
346681
Forward Design 1
score (lower is better): 2
                                                 AAATTAATACGACTCACTAT
:back translated gRNA
                                 ATCGATCTCGATCCCGCGAAATTAATACGACTCACTA-----GAATTGTGAGCGGATAACAATTTCAGAATTCAAAAGA
:repair template homology
5':GCCACGATGCGTCCGGCGTAGAGGATCGAGATCGATCTCGATCCCGCGAAATTAATACGACTCACTATAGGGGAATTGTGAGCGGATAACAATTTCAGAATTCAAAAGATCTTT
TAAGAAGGAGATATAC:3'
3':CGGTGCTACGCAGGCCGCATCTCCTAGCTCTAGCTAGAGCTAGGGCGCTTTAATTATGCTGAGTGATATCCCCTTAACACTCGCCTATTGTTAAAGTCTTAAGTTTTCTAGAAA
ATTCTTCCTCTATATG:5'

Repair Template:
5':ATCGATCTCGATCCCGCGAAATTAATACGACTCACTATAGAGGGAATTGTGAGCGGATAACAATTTCAGAATTCAAAAGA:3'

346681
Forward Design 2
score (lower is better): 2
                                                               CACTATAGGGGAATTGTGAG
:back translated gRNA
                                 ATCGATCTCGATCCCGCGAAATTAATACGACTCACTA-----GAATTGTGAGCGGATAACAATTTCAGAATTCAAAAGA
:repair template homology
5':GCCACGATGCGTCCGGCGTAGAGGATCGAGATCGATCTCGATCCCGCGAAATTAATACGACTCACTATAGGGGAATTGTGAGCGGATAACAATTTCAGAATTCAAAAGATCTTT
TAAGAAGGAGATATAC:3'
3':CGGTGCTACGCAGGCCGCATCTCCTAGCTCTAGCTAGAGCTAGGGCGCTTTAATTATGCTGAGTGATATCCCCTTAACACTCGCCTATTGTTAAAGTCTTAAGTTTTCTAGAAA
ATTCTTCCTCTATATG:5'

Repair Template:
5':ATCGATCTCGATCCCGCGAAATTAATACGACTCACTATAGAGGGAATTGTGAGCGGATAACAATTTCAGAATTCAAAAGA:3'

4279280
Reverse Design 1
score (lower is better): 2
3':CATATAGAGGAAGAATTTTCTAGAAAACTTAAGACTTTAACAATAGGCGAGTGTTAAGGGGATATCACTCAGCATAATTAAAGCGCCCTAGCTCTAGCTAGAGCTAGGAGATGC
GGCCTGCGTAGCACCG:5'
5':GTATATCTCCTTCTTAAAAGATCTTTTGAATTCTGAAATTGTTATCCGCTCACAATTCCCCTATAGTGAGTCGTATTAATTTCGCGGGATCGAGATCGATCTCGATCCTCTACG
CCGGACGCATCGTGGC:3'
                      GATCTTTTGAATTCTGAAATTGTTATCCGCTCACAAT---CCTATAGTGAGTCGTATTAATTTCGCGGGATCGAGAT
:repair template homology
                                                                   AGTGAGTCGTATTAATTTCG
:back translated gRNA

Repair Template:
5':GATCTTTTGAATTCTGAAATTGTTATCCGCTCACAATTCCTCCCTATAGTGAGTCGTATTAATTTCGCGGGATCGAGAT:3'

bpsA
346681
Forward Design 1
score (lower is better): 2
                                                 AAATTAATACGACTCACTAT
:back translated gRNA
                                 ATCGATCTCGATCCCGCGAAATTAATACGACTCACTA-----GAATTGTGAGCGGATAACAATTTCAGAATTCAAAAGA
:repair template homology
5':GCCACGATGCGTCCGGCGTAGAGGATCGAGATCGATCTCGATCCCGCGAAATTAATACGACTCACTATAGGGGAATTGTGAGCGGATAACAATTTCAGAATTCAAAAGATCTTT
TAAGAAGGAGATATAC:3'
3':CGGTGCTACGCAGGCCGCATCTCCTAGCTCTAGCTAGAGCTAGGGCGCTTTAATTATGCTGAGTGATATCCCCTTAACACTCGCCTATTGTTAAAGTCTTAAGTTTTCTAGAAA
ATTCTTCCTCTATATG:5'

Repair Template:
5':ATCGATCTCGATCCCGCGAAATTAATACGACTCACTATAGAGGGAATTGTGAGCGGATAACAATTTCAGAATTCAAAAGA:3'

346681
Forward Design 2
score (lower is better): 2
                                                               CACTATAGGGGAATTGTGAG
:back translated gRNA
                                 ATCGATCTCGATCCCGCGAAATTAATACGACTCACTA-----GAATTGTGAGCGGATAACAATTTCAGAATTCAAAAGA
:repair template homology
5':GCCACGATGCGTCCGGCGTAGAGGATCGAGATCGATCTCGATCCCGCGAAATTAATACGACTCACTATAGGGGAATTGTGAGCGGATAACAATTTCAGAATTCAAAAGATCTTT
TAAGAAGGAGATATAC:3'
3':CGGTGCTACGCAGGCCGCATCTCCTAGCTCTAGCTAGAGCTAGGGCGCTTTAATTATGCTGAGTGATATCCCCTTAACACTCGCCTATTGTTAAAGTCTTAAGTTTTCTAGAAA
ATTCTTCCTCTATATG:5'

Repair Template:
5':ATCGATCTCGATCCCGCGAAATTAATACGACTCACTATAGAGGGAATTGTGAGCGGATAACAATTTCAGAATTCAAAAGA:3'

4279280
Reverse Design 1
score (lower is better): 2
3':CATATAGAGGAAGAATTTTCTAGAAAACTTAAGACTTTAACAATAGGCGAGTGTTAAGGGGATATCACTCAGCATAATTAAAGCGCCCTAGCTCTAGCTAGAGCTAGGAGATGC
GGCCTGCGTAGCACCG:5'
5':GTATATCTCCTTCTTAAAAGATCTTTTGAATTCTGAAATTGTTATCCGCTCACAATTCCCCTATAGTGAGTCGTATTAATTTCGCGGGATCGAGATCGATCTCGATCCTCTACG
CCGGACGCATCGTGGC:3'
                      GATCTTTTGAATTCTGAAATTGTTATCCGCTCACAAT---CCTATAGTGAGTCGTATTAATTTCGCGGGATCGAGAT
:repair template homology
                                                                   AGTGAGTCGTATTAATTTCG
:back translated gRNA

Repair Template:
5':GATCTTTTGAATTCTGAAATTGTTATCCGCTCACAATTCCTCCCTATAGTGAGTCGTATTAATTTCGCGGGATCGAGAT:3'
```

In [24]:
```python
#Other Promoter To Alter

#Information about it here: https://biocyc.org/ECOLI/NEW-IMAGE?type=OPERON&object=TU0-5901
promoter = 'gcataaaaagggttatccaaaggtcattgcaccaacatggtgcttaatgtttccattgaagcactatattggtgcaacattcaca'

#Create List of outseqs
outseq =   'gcataaaaagggttatccaaaggCcattgcaccaacatggtgcttaatgtttccattgaagcactatattggtgcaacattcaca'

#Create List of Designs
in_seq  = Seq(promoter.upper(),DNAAlphabet())
out_seq = Seq(outseq.upper(),DNAAlphabet())

designs = mage_design(jbei_19353,in_seq.reverse_complement(),out_seq.reverse_complement(),max_template_length=80)

for i,design in enumerate(designs):
    genome = jbei_19353[(design[-1]-30):(design[-1]+100)]
    print(design[-1])
    print('Forward Design {}'.format(i+1))
    print('score (lower is better): {}'.format(score_gRNA(jbei_19353,design[0])))
    print_design(genome,design[0],designs[0][1],ha_length=37)
    print('')

designs = mage_design(jbei_19353.reverse_complement(),in_seq,out_seq,max_template_length=80)
for i,design in enumerate(designs):
    genome = jbei_19353.reverse_complement()[(design[-1]-30):(design[-1]+100)]
    #print(design)
    print(design[-1])
    print('Reverse Design {}'.format(i+1))
    print('score (lower is better): {}'.format(score_gRNA(jbei_19353,design[0])))
    print_design(genome,design[0],designs[0][1],forward=False,ha_length=35)
    print('')
```

```
4053532
Forward Design 1
score (lower is better): 1
                     GCACCAATATAGTGCTTCAA
:back translated gRNA
                                 TGCTTCAATGGAAACATTAAGCACCATGTTGGTGCAA-----TTTGGATAACCCTTTTTATGCTCCGTGAAAGCGATCA
:repair template homology
5':CACCACGATGTGAATGTTGCACCAATATAGTGCTTCAATGGAAACATTAAGCACCATGTTGGTGCAATGACCTTTGGATAACCCTTTTTATGCTCCGTGAAAGCGATCACAAAG
GGACTCTGCAATACTT:3'
3':GTGGTGCTACACTTACAACGTGGTTATATCACGAAGTTACCTTTGTAATTCGTGGTACAACCACGTTACTGGAAACCTATTGGGAAAAATACGAGGCACTTTCGCTAGTGTTTC
CCTGAGACGTTATGAA:5'

Repair Template:
5':TGCTTCAATGGAAACATTAAGCACCATGTTGGTGCAATGGCCTTTGGATAACCCTTTTTATGCTCCGTGAAAGCGATCA:3'

4053532
Forward Design 2
score (lower is better): 1
                                          GGAAACATTAAGCACCATGT
:back translated gRNA
                                 TGCTTCAATGGAAACATTAAGCACCATGTTGGTGCAA-----TTTGGATAACCCTTTTTATGCTCCGTGAAAGCGATCA
:repair template homology
5':CACCACGATGTGAATGTTGCACCAATATAGTGCTTCAATGGAAACATTAAGCACCATGTTGGTGCAATGACCTTTGGATAACCCTTTTTATGCTCCGTGAAAGCGATCACAAAG
GGACTCTGCAATACTT:3'
3':GTGGTGCTACACTTACAACGTGGTTATATCACGAAGTTACCTTTGTAATTCGTGGTACAACCACGTTACTGGAAACCTATTGGGAAAAATACGAGGCACTTTCGCTAGTGTTTC
CCTGAGACGTTATGAA:5'

Repair Template:
5':TGCTTCAATGGAAACATTAAGCACCATGTTGGTGCAATGGCCTTTGGATAACCCTTTTTATGCTCCGTGAAAGCGATCA:3'

4053532
Forward Design 3
score (lower is better): 1
                                                         CATGTTGGTGCAATGACCTT
:back translated gRNA
                                 TGCTTCAATGGAAACATTAAGCACCATGTTGGTGCAA-----TTTGGATAACCCTTTTTATGCTCCGTGAAAGCGATCA
:repair template homology
5':CACCACGATGTGAATGTTGCACCAATATAGTGCTTCAATGGAAACATTAAGCACCATGTTGGTGCAATGACCTTTGGATAACCCTTTTTATGCTCCGTGAAAGCGATCACAAAG
GGACTCTGCAATACTT:3'
3':GTGGTGCTACACTTACAACGTGGTTATATCACGAAGTTACCTTTGTAATTCGTGGTACAACCACGTTACTGGAAACCTATTGGGAAAAATACGAGGCACTTTCGCTAGTGTTTC
CCTGAGACGTTATGAA:5'

Repair Template:
5':TGCTTCAATGGAAACATTAAGCACCATGTTGGTGCAATGGCCTTTGGATAACCCTTTTTATGCTCCGTGAAAGCGATCA:3'

583354
Reverse Design 1
score (lower is better): 1
3':AGGCGTTTGTTCATAACGTCTCAGGGAAACACTAGCGAAAGTGCCTCGTATTTTTCCCAATAGGTTTCCAGTAACGTGGTTGTACCACGAATTACAAAGGTAACTTCGTGATAT
AACCACGTTGTAAGTG:5'
5':TCCGCAAACAAGTATTGCAGAGTCCCTTTGTGATCGCTTTCACGGAGCATAAAAAGGGTTATCCAAAGGTCATTGCACCAACATGGTGCTTAATGTTTCCATTGAAGCACTATA
TTGGTGCAACATTCAC:3'
                                 TGATCGCTTTCACGGAGCATAAAAAGGGTTATCCA---------GCACCAACATGGTGCTTAATGTTTCCATTGAAGCA
:repair template homology
                         TCCCTTTGTGATCGCTTTCA
:back translated gRNA

Repair Template:
5':TGATCGCTTTCACGGAGCATAAAAAGGGTTATCCAAAGGCCCATTGCACCAACATGGTGCTTAATGTTTCCATTGAAGCA:3'

583354
Reverse Design 2
score (lower is better): 1
3':AGGCGTTTGTTCATAACGTCTCAGGGAAACACTAGCGAAAGTGCCTCGTATTTTTCCCAATAGGTTTCCAGTAACGTGGTTGTACCACGAATTACAAAGGTAACTTCGTGATAT
AACCACGTTGTAAGTG:5'
5':TCCGCAAACAAGTATTGCAGAGTCCCTTTGTGATCGCTTTCACGGAGCATAAAAAGGGTTATCCAAAGGTCATTGCACCAACATGGTGCTTAATGTTTCCATTGAAGCACTATA
TTGGTGCAACATTCAC:3'
                                 TGATCGCTTTCACGGAGCATAAAAAGGGTTATCCA---------GCACCAACATGGTGCTTAATGTTTCCATTGAAGCA
:repair template homology
                                     CGCTTTCACGGAGCATAAAA
:back translated gRNA

Repair Template:
5':TGATCGCTTTCACGGAGCATAAAAAGGGTTATCCAAAGGCCCATTGCACCAACATGGTGCTTAATGTTTCCATTGAAGCA:3'

583354
Reverse Design 3
score (lower is better): 1
3':AGGCGTTTGTTCATAACGTCTCAGGGAAACACTAGCGAAAGTGCCTCGTATTTTTCCCAATAGGTTTCCAGTAACGTGGTTGTACCACGAATTACAAAGGTAACTTCGTGATAT
AACCACGTTGTAAGTG:5'
5':TCCGCAAACAAGTATTGCAGAGTCCCTTTGTGATCGCTTTCACGGAGCATAAAAAGGGTTATCCAAAGGTCATTGCACCAACATGGTGCTTAATGTTTCCATTGAAGCACTATA
TTGGTGCAACATTCAC:3'
                                 TGATCGCTTTCACGGAGCATAAAAAGGGTTATCCA---------GCACCAACATGGTGCTTAATGTTTCCATTGAAGCA
:repair template homology
                                                 GCATAAAAAGGGTTATCCAA
:back translated gRNA

Repair Template:
5':TGATCGCTTTCACGGAGCATAAAAAGGGTTATCCAAAGGCCCATTGCACCAACATGGTGCTTAATGTTTCCATTGAAGCA:3'

583354
Reverse Design 4
score (lower is better): 1
3':AGGCGTTTGTTCATAACGTCTCAGGGAAACACTAGCGAAAGTGCCTCGTATTTTTCCCAATAGGTTTCCAGTAACGTGGTTGTACCACGAATTACAAAGGTAACTTCGTGATAT
AACCACGTTGTAAGTG:5'
5':TCCGCAAACAAGTATTGCAGAGTCCCTTTGTGATCGCTTTCACGGAGCATAAAAAGGGTTATCCAAAGGTCATTGCACCAACATGGTGCTTAATGTTTCCATTGAAGCACTATA
TTGGTGCAACATTCAC:3'
                                 TGATCGCTTTCACGGAGCATAAAAAGGGTTATCCA---------GCACCAACATGGTGCTTAATGTTTCCATTGAAGCA
:repair template homology
                                                                  CAAAGGTCATTGCACCAACA
:back translated gRNA

Repair Template:
5':TGATCGCTTTCACGGAGCATAAAAAGGGTTATCCAAAGGCCCATTGCACCAACATGGTGCTTAATGTTTCCATTGAAGCA:3'
```

In [15]:
```python
#Pick Good Designs
designs = [['sfp' ,'AAATTAATACGACTCACTAT','ATCGATCTCGATCCCGCGAAATTAATACGACTCACTATAGAGGAATTGTGAGCGGATAACAATTTCAGAATTCAA
AAGA'],   #Pam Mutation in Active Region
           ['sfp' ,'CACTATAGGGGAATTGTGAG','ATCGATCTCGATCCCGCGAAATTAATACGACTCACTATAGAGGAATTGTGAGCGAATAACAATTTCAGAATTCAA
AAGA'],
           ['glnA','TCCCTTTGTGATCGCTTTCA','TGATCGCTTTCACGAAGCATAAAAAGGGTTATCCAAAGGTCACTGCACCAACATGGTGCTTAATGTTTCCATTGA
AGCA'],
           ['glnA','CGCTTTCACGGAGCATAAAA','TGATCGCTTTCACGGAGCATAAAAAGAGTTATCCAAAGGTCGTTGCACCAACATGGTGCTTAATGTTTCCATTGA
AGCA'],
           ['glnA','GCATAAAAAGGGTTATCCAA','TGATCGCTTTCACGGAGCATAAAAAGGGTTATCCAAAGTTCATTGCACCAACATGGTGCTTAATGTTTCCATTGA
AGCA'],   #Pam Mutation in Active Region
           ['glnA','CAAAGGTCATTGCACCAACA','TGATCGCTTTCACGGAGCATAAAAAGGGTTATCCAAAGGCCATTGCACCAACATGATGCTTAATGTTTCCATTGA
AGCA']]

# 'ag:gtcatt:' -> Mutation Region
```

In [25]:
```python
rep_temp = 'ATCGATCTCGATCCCGCGAAATTAATACGACTCACTATAGAGGAATTGTGAGCGGATAACAATTTCAGAATTCAAAAGA'
print(len(rep_temp))
```

```
79
```

In [ ]:
